# Fast, lightweight, and accurate metagenomic functional profiling using FracMinHash sketches

**DOI:** 10.1101/2023.11.06.565843

**Authors:** Mahmudur Rahman Hera, Shaopeng Liu, Wei Wei, Judith S. Rodriguez, Chunyu Ma, David Koslicki

**Author notes:** The authors contributed equally to the work (Mahmudur Rahman Hera, Shaopeng Liu).

## Abstract

**Motivation:** Functional profiling of metagenomic samples is essential to decipher the functional capabilities of microbial communities. Traditional and more widely used functional profilers in the context of metagenomics rely on aligning reads against a known reference database. However, aligning sequencing reads against a large and fast-growing database is computationally expensive. In general, *k*-mer-based sketching techniques have been successfully used in metagenomics to address this bottleneck, notably in taxonomic profiling. In this work, we describe leveraging FracMinHash (implemented in sourmash, a publicly available software), a *k*-mer-sketching algorithm, to obtain functional profiles of metagenome samples.

**Results:** We show how pieces of the `sourmash` software (and the resulting FracMinHash sketches) can be put together in a pipeline to functionally profile a metagenomic sample. We named our pipeline `fmh-funprofiler`. We report that the functional profiles obtained using this pipeline demonstrate comparable completeness and better purity compared to the profiles obtained using other alignment-based methods when applied to simulated metagenomic data. We also report that `fmh-funprofiler` is 39-99x faster in wall-clock time, and consumes up to 40-55x less memory. Coupled with the KEGG database, this method not only replicates fundamental biological insights but also highlights novel signals from the Human Microbiome Project datasets.

**Reproducibility:** This fast and lightweight metagenomic functional profiler is freely available and can be accessed here: https://github.com/KoslickiLab/fmh-funprofiler. All scripts of the analyses we present in this manuscript can be found on [GitHub](https://github.com/KoslickiLab/KEGG_sketching_annotation_reproducibles).

## 1 Introduction

Metagenomic profiling, the process of evaluating and identifying the genomic information present within a given environmental sample, offers valuable insights into the genetic diversity, functional capacity, and ecological contributions of microorganisms. Specifically, metagenomic functional profiling (a.k.a. functional annotation), the computational process that involves identifying and quantifying functional components within metagenomic data, is essential to understanding microbial functionality, phenotype-genotype association, host interaction, disease progress, etc. [13, 14, 44, 58, 47].

A related and frequently examined entity in biology, especially in gene studies, is orthologous genes. If two genes in two different species are evolutionary descendants of the same gene in the least common ancestor of these species, then these two genes are called orthologs [62]. The study of orthologous genes contributes crucially to comparative genomics and evolutionary studies, and often these studies make use of the well-known fact that orthologous genes often carry out equivalent or identical biological functions across species boundaries [20, 39, 64]. For example, researchers have successfully identified functions of genes in newly sequenced genomes using orthology in model organisms [62, 39, 10]. Previously, it was common practice to annotate the function of human genes using the orthologs in mice, as it is easier to experiment on the latter [25]. Largely speaking, orthology traces genes to the same ancestral gene, and therefore, orthologous relationships are considered to be the most accurate way to capture differences and similarities in the composition of genomes from different species [18]. Naturally, computational efforts have explored the area, and constructed databases of orthologous genes, such as AspGD [3], MBGD [70], OrthoDB [41], and KEGG [34, 33]. Coming back to the study of genes in metagenomes: it is crucial to note the numerous computational tools that have been developed to identify genes in an environmental sample. In a broader sense, these tools fall into two categories: (a) gene prediction tools – which attempt to predict genes *ab initio* using only the sequences [50, 61, 80], and (b) gene classification/annotation tools – which aim to annotate a metagenome sample against a database of known genes. The latter category is more relevant for functional profiling since we need to have prior knowledge of the functions of known genes. The majority of the tools in this category rely on alignment-based approaches. For example, `eggNOG-Mapper` performs alignments with orthologs utilizing profile HMM models for query assignment [27]. `MG-RAST` [37] uses `BLAST` [77] to search against the M5nr database [73]. `BlastKOALA` [35], `GhostKOALA` [35], and `KofamSCAN` [2], which utilize the BLASTP [77], GHOSTX [66], and hmmsearch [32] algorithms respectively, are developed to search against Kyoto Encyclopedia of Genes and Genomes (KEGG) database [34]. Perhaps the fastest (and most widely used) tool in the context is `DIAMOND` [7], which utilizes double indexing on both query and reference to enable fast alignment. `DIAMOND’s` novel algorithm can match the sensitivity of `BLAST` while having much better efficiency [7]. A couple of years after DIAMOND was released, MMSeqs2 was published as another fast and sensitive protein aligner tool [63] and has been very widely used in massive data processing pipelines, although recent benchmarking studies have shown that DIAMOND runs faster than MMSeqs2 in finding orthologs as reciprocal best hits [22].

Despite the popular use of these tools, the primary use of alignment-based algorithms makes these a poor practical choice in terms of scalability. With the ever-growing volume of sequencing data, alignment-based methods, despite their historical success, will eventually be overshadowed by more scalable techniques. Computational biologists, therefore, continue to turn to sketching-based methods, which are often faster and more lightweight; and theoretical guarantees of the sketching algorithms ensure their high accuracy. These alignment-based tools also lack the use of orthology relationships of the genes. Using well-studied orthologous groups has the potential to make functional profiling pipelines more efficient, both in time and memory.

In this paper, we present a sketching-based pipeline to functionally profile a metagenome in terms of orthologous gene groups. We used FracMinHash as our sketching technique to develop the pipeline – recent investigations have shown the successful use of FracMinHash (used in the software package `sourmash` [6]) in metagenomics analysis [49]. We used KEGG as the database of orthologous genes. The orthologous groups in KEGG are called KOs. Our pipeline can take FracMinHash sketches of a given metagenome and the reference KOs, and discover what KOs are present in the metagenome. The pipeline can also annotate the relative abundances of the KOs. It is fast and lightweight because of using FracMinHash sketches and is nearly as accurate as alignment-based tools. To critically assess if and when sketching should be used over alignment for metagenomic functional profiling, we investigated the performance and resource usage of the pipeline against popular protein alignment tool `DIAMOND`. We also used the pipeline to functionally profile Human Gut Microbiota. Furthermore, the KO groups are very well studied in the KEGG database in the form of pathways – which allowed us to make insightful deductions about these microbiota. The pipeline is freely available and can be accessed here: https://github.com/KoslickiLab/fmh-funprofiler.

## 2 Background

### 2.1 KEGG and gene orthology

Selecting an appropriate reference database is crucial in metagenomic functional profiling. Orthologous genes, which share a common evolutionary ancestry, are commonly employed for inferring functions [12]. Numerous efforts have been undertaken to construct databases for gene orthology, such as NCBI COG [19], AGNOSTOS-DB [72], OrthoDB [41], and Ensembl Compara [23]. The KEGG database is a comprehensive and integrated resource for biological interpretation by providing gene annotations and mapping genes to manually created pathway maps [34, 33]. These KO groups serve as a foundation for functional annotation in metagenomics and can be further connected into pathway maps to illustrate the hierarchical interconnections of biological functions within living organisms [35, 42, 15].

### 2.2 FracMinHash and sketching-based methods

With a large number of reference genomes and an exponential surge in metagenomic data production, there is an imperative need for the creation of computational models that are both scalable and robust, ensuring precision in analysis. *K*-mer-based algorithms, particularly sketching-based methods, are gaining increasing popularity for metagenomic profiling implementations in this regard. A *k*-mer is a sequence of *k* consecutive nucleotides, extracted from a longer sequence. Algorithms designed to work with *k*-mers split the entire sample into *k*-mers, and analyze the number of shared/dissimilar *k*-mers among multiple samples. The number of all distinct *k*-mers in a sequencing sample can often be huge, so sketching-based methods take a fingerprint of the *k*-mers (called a *sketch*) and work with these much smaller sets, ensuring less consumption of computational resources. The most popular sketching method for many years has been MinHash, introduced in the context of document comparisons [5]. Mash [53] was developed to apply MinHash to genomic data and has been very widely used. However, recent studies have shown that the error in comparing two MinHash sketches depends on the sketch size [56], and the size needs to grow quadratically to compensate for the error [53, Fig S1]. More recent works have shown that when sets of very dissimilar sizes are compared, MinHash sketches perform relatively poorly [45, 40]. Researchers have proposed many adjustments to MinHash to tackle this issue [52, 40, 4, 30]; using a variable sketch size (instead of MinHash’s fixed-size scheme), the recently introduced FracMinHash sketch [29, 21] is one such example. Simply speaking, a FracMinHash sketch retains a fraction of the *k*-mers in the original sample. Formally, given a perfect hash function *h* : Ω → [0, *H*] for some *H* ∈ ℝ and a *scale factor s* where 0 ≤ *s* ≤ 1, a FracMinHash sketch of a set *A* is defined as follows:

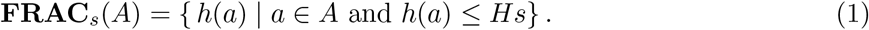

The scale factor *s* is a tunable parameter that can modify the size of the sketch. For a fixed *s*, if the set *A* grows larger, the sketch **FRAC**_*s*_(*A*) grows proportionally. This sketching technique was first introduced in the software package `sourmash` [6, 55], and has recently been theoretically analyzed to justify its use [21]. `sourmash` has successfully used FracMinHash to obtain genome-wide comparisons of biological sequences. In the second round of Critical Assessment of Metagenomic Interpretation (CAMI) challenges [59, 49], the tool `sourmash gather` [28], exhibited the highest completeness and purity in taxonomic profiling from metagenomic data for multiple datasets [49] across the genus and the species levels.

## 3 Results

We designed a *k*-mer and sketching-based pipeline (shown in Figure 1) to identify orthologs in a metagenome. `fmh-funprofiler` computes FracMinHash sketches of the orthologous gene groups in a reference database. It then computes the FracMinHash sketch of an input metagenome and queries it against the reference sketches to detect which orthologous gene groups are present in the metagenome. The entire pipeline is described in more detail in the Methods section. Ideally, this pipeline works with any ortholog database, such as OrthoDB or KEGG. In this work, we used KEGG as the ortholog database and set `fmh-funprofiler` to identify KEGG Orthologs (KOs) in metagenomes. A downsampling factor of 1000 was used in every analysis – which is the default in the software package `sourmash`.

**Figure 1:**
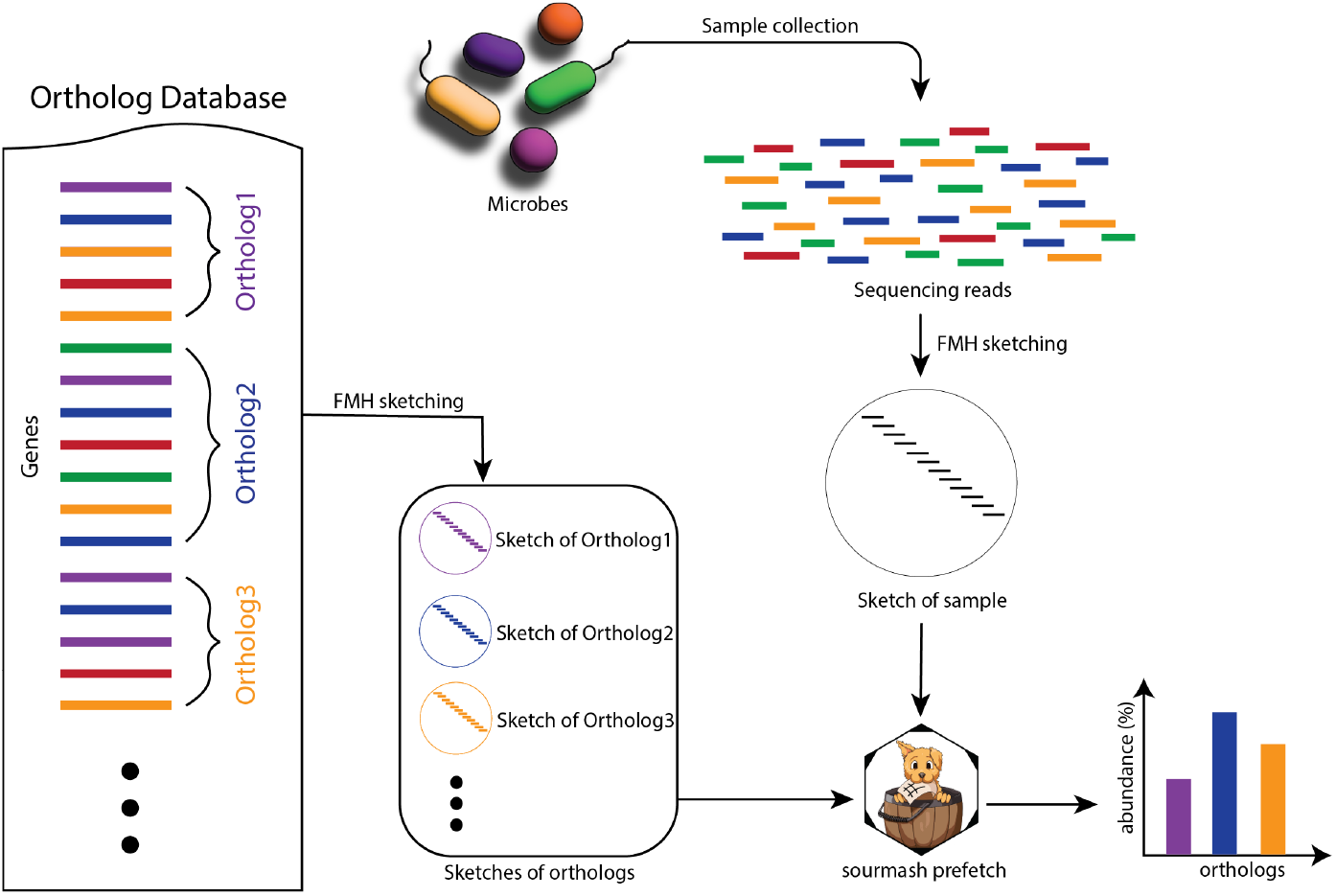
`fmh-funprofiler` identifies orthologs in a metagenome by splitting orthologs into *k*-mers and creating FracMinHash sketches with `sourmash sketch`. It then sketches the input metagenome file in the same way and computes the sample composition using the orthologs’ *k*-mers with `sourmash prefetch`. Finally, it post-processes the output to compute the abundances.

### 3.1 fmh-funprofiler reveals orthologs accurately

We start presenting the results by benchmarking the performance of `fmh-funprofiler` in identifying KOs in metagenomes. We used CAMISIM [16] to simulate the metagenomes using a random selection of all bacterial genomes present in the KEGG database, as of June 2023. From the output of CAMISIM, we identified the KOs truly present in the simulation along with their relative abundances. Details of these simulations and finding the ground truth are elaborated in the Methods section.

We made comprehensive comparisons with `DIAMOND` [7], which is the fastest and most widely used protein alignment tool for metagenomic data, reported to be more than 20,000 times faster than BLASTX [7]. We also obtained the HMM profiles of all the KEGG Orthologs published by KEGG and assigned KOs to simulated metagenomes using KEGG’s tool `KofamScan` [2]. The results using `KofamScan` are not included here because (a) we found that `KofamScan` requires unrealistic computational resources (requires about seven days to complete for a metagenome with 1M reads), and (b) `KofamScan` is designed to run on a small number of assembled genomes, not a huge number of short reads. Despite being fast and sensitive, we did not include `MMSeqs2` in our experiments because benchmarking studies have shown that `DIAMOND` runs faster than `MMSeqs2` in “fast” mode, and is more sensitive in “sensitive” mode [22].

To run `DIAMOND`, we used the protein sequences of all the genes in the KEGG database as the references. As an output, `DIAMOND` matches the reads in the metagenome to possible genes. `DIAMOND` reports multiple alignments, and we filtered the best match to find the gene assigned to a read. We also recorded the number of nucleotides matched with the target gene. We then used the gene-to-KO grouping and the number of overlapping nucleotides to compute the relative abundances of the KOs. We used 128 threads to invoke each run of `DIAMOND`. We also ran `DIAMOND` in both “fast” and “sensitive” modes. We found that the “fast” mode is already sensitive enough for metagenomic functional profiling (identifies almost all preset KOs), and thus the “sensitive” mode ends up consuming more resources with no additional benefit. Therefore, we are only showing comparisons of our pipeline against `DIAMOND fast`. To make these comparisons, we computed the following performance metrics: precision, completeness, weighted Jaccard similarity, Pearson correlation coefficient, and Bray-Curtis distance. For the first four metrics, higher implies better. For Bray-Curtis, lower implies better. For the sake of brevity, we have included elaborate definitions of these metrics in the supplemental materials.

We compared the performance of `DIAMOND` and `fmh-funprofiler` using simulated metagenomes across different factors: by varying the size of the metagenomes, the number of organisms (both real and novel) in the metagenomes, the rate of divergence of novel strains, and the rate of error of the sequencing technology. In all these studies, we found consistent results. For the sake of brevity, we are only showing the performance metrics for various rates of error of the sequencing technology in Figure 2. In this comparison, we simulated metagenomes of size 64Gbp using CAMISIM, used `wgsim` for read generation, and varied the rate of sequencing error. For every error rate, we simulated 30 metagenomes using different seeds. Figure 2 shows the mean performance metrics, where the error bars indicate one standard deviation.

**Figure 2:**
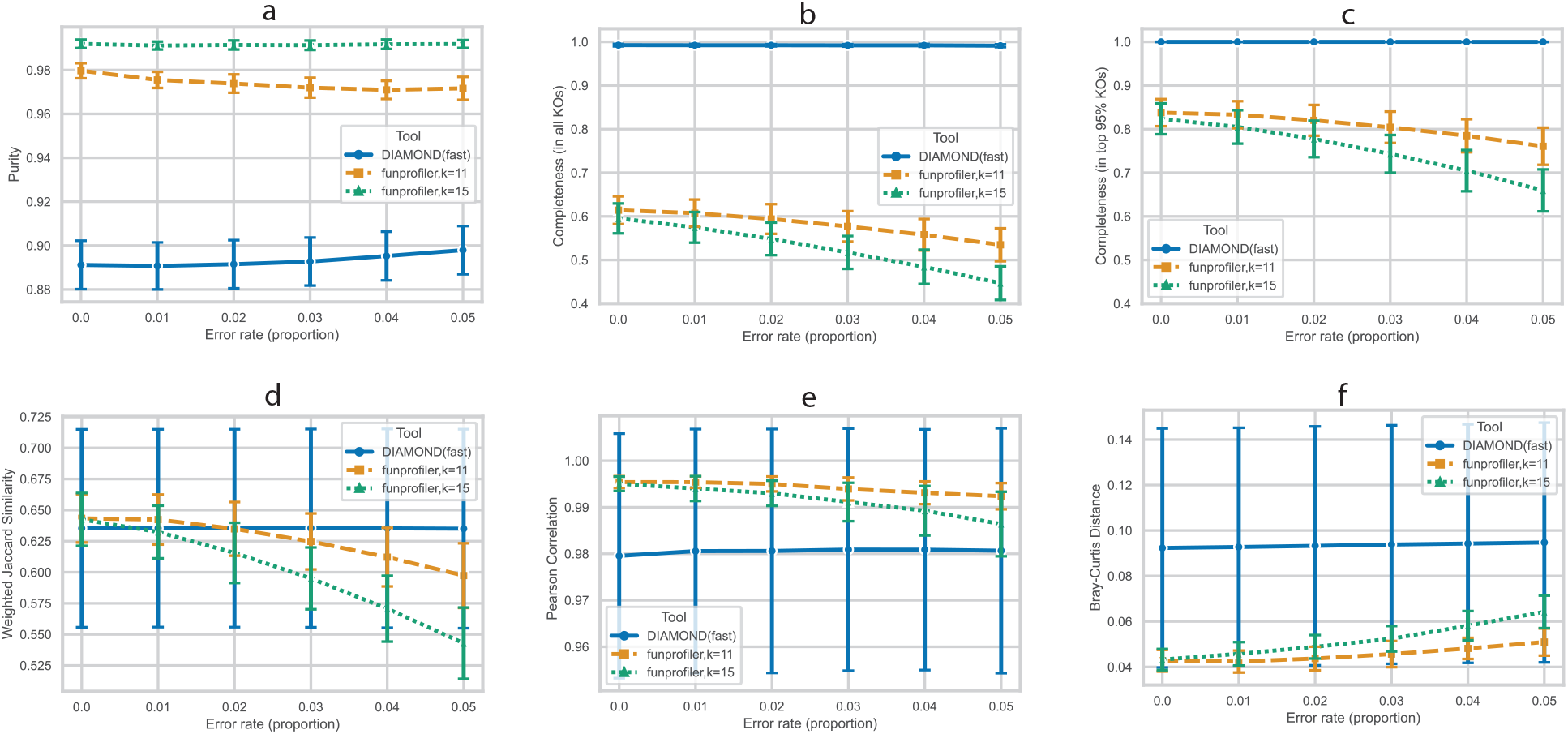
Average performances in identifying KOs in simulated metagenomes. The metagenomes are of size 6.4 Gbp and consist of 6.1-6.9 thousand KOs. Metrics shown: (a) purity, (b) completeness in all KOs, (c) completeness in top 95% abundant KOs, (d) weighted Jaccard index between ground truth KOs and identified KOs, (e) Pearson correlation coefficient of relative abundances computed by the tools and the ground truth, and (f) Bray-Curtis distance of the relative abundances computed by the tools and the ground truth. Every point shows an average over 30 random seeds (30 different simulations), and error bars indicate one standard deviation.

The comparisons show that `fmh-funprofiler` is very precise with few false positives – most KOs identified by `fmh-funprofiler` are in the ground truth. `DIAMOND`, on the other hand, detects some false positives, but also at the same time, is extremely sensitive – discovering *all* ground truth KOs in every setting. In contrast, `fmh-funprofiler` exhibits low completeness. This is expected, given the lossy downsampling nature of the sketching algorithm. The *k*-mers in the KOs that are very lowly abundant are expected to not be included in the FracMinHash sketches. Consequently, the absence of lowly-abundant KOs in the sketches results in low completeness. If we restrict to the KOs that constitute the top 95% of the abundances in the ground truth (by excluding the very lowly abundant KOs), the completeness of `fmh-funprofiler` improves to up to 85% as seen in panel c of 2.

It is important to note that there is a trade-off between purity and completeness. `DIAMOND`, although detects all KOs, ends up detecting some false positives. `fmh-funprofiler` detects a fewer number of KOs more precisely. To get a holistic idea, we computed the Jaccard index of the predicted KO abundances and the ground truth KO abundances. This shows that `fmh-funprofiler` is comparable to the alignment-based tool `DIAMOND`. Additionally, to investigate the KO abundances computed by the tools, we analyzed correlation and distance with the ground truth. Our analysis found that the abundances computed by `fmh-funprofiler` correlate with the ground truth more closely compared to `DIAMOND`, with a smaller Bray-Curtis distance.

### 3.2 fmh-funprofiler uses less computational resources

We also recorded time and memory to process all 30 metagenomes using these tools. `DIAMOND` was run using 128 threads. `fmh-funprofiler` itself is single-threaded, and so we ran `fmh-funprofiler` on 30 threads, one for every metagenome. Wall-clock time and peak memory usage are shown in Figure 3. It shows that `fmh-funprofiler` can finish in one 39th to one 99th fraction of the time required by `DIAMOND`. Note that `DIAMOND` was run in `fast` mode. We also found that `fmh-funprofiler` requires 40-55x less memory to operate.

**Figure 3:**
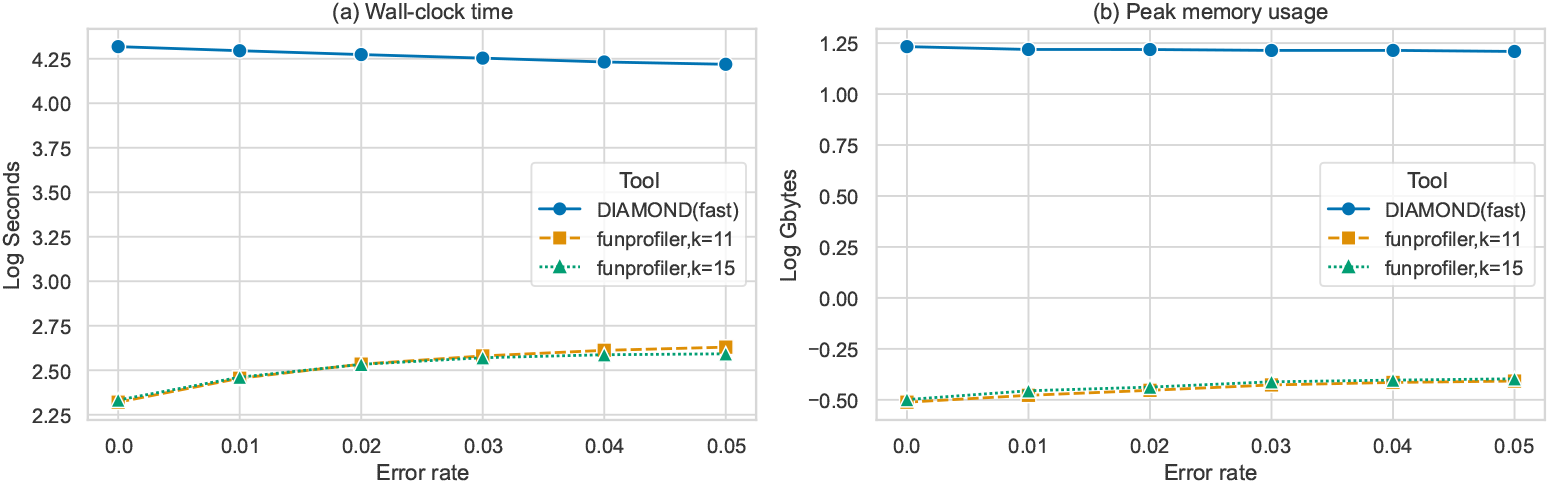
Computational resources consumed by `DIAMOND` and `fmh-funprofiler` in identifying KOs in 30 simulated metagenomes: (a) shows total wall-clock time, and (b) shows peak memory usage. The metagenomes are the same ones used to generate Figure 2. `DIAMOND` was run using 128 threads, and `fmh-funprofiler` using only 30 (one for each input). We found that `fmh-funprofiler` runs 39-99x faster, and uses 40-55x less memory compared to `DIAMOND`.

The results shown in Figures 2 and 3 indicate the following: `fmh-funprofiler` is expected to miss low abundance KOs, but detects the more abundant KOs, is extremely precise, and the relative abundances computed by `fmh-funprofiler` are more accurate at smaller error rates. Besides being comparable in performance, `fmh-funprofiler` is also fast and lightweight compared to alignment-based tools.

### 3.3 fmh-funprofiler reveals functional insights of human metagenomic data

While the accuracy and improved efficiency of `fmh-funprofiler` have been demonstrated in simulated data, we subsequently employ this approach on actual human metagenomic data from the Human Microbiome Project (HMP)[69]. We use the KEGG (Kyoto Encyclopedia of Genes and Genomes) database[34], with KEGG Orthology (KOs) specifically, to profile metagenomic functions. The analysis encompasses a total of 1747 filtered samples, spanning 6 ethnic groups, 2 genders, and 3 study groups.

Functional profiles reveal the presence of housekeeping KOs and pathways consistently observed across the majority of samples, irrespective of conditions (Supplementary Figure 5). In addition to the core housekeeping KOs present in most samples, numerous other KOs appear in just one or two conditions. The top hits of most observed KOs include K01872 for Aminoacyl-tRNA biosynthesis, K03701 for excinuclease ABC subunit A, and K02004 for (putative) ABC transport system permease protein. Most of the top hits are recognized as vital cellular functions, thus constituting a compilation of housekeeping functions. More universal functional insights of human microbiota can be obtained from the right tail of Supplementary Figure 5b. We further checked the functional pathways at KEGG level 2 and level 3 and found that genes in Biosynthesis of amino acids (Ko01230), Carbon metabolism (Ko01200), and Amino sugar and nucleotide sugar metabolism (Ko00520) are abundant in most samples. These observations are consistent with the previous functional analysis in human gut microbiomes [82, 43, 68, 54], and suggest that these, along with Purine metabolism (Ko00230), Amino sugar and nucleotide sugar metabolism (Ko00520), and Glycolysis / Gluconeogenesis (Ko00010) etc., are the core functions of the gut microbiota.

Next, we analyze the distinct functions among different conditions (Type 2 Diabetes, T2D; Healthy, HHS; and Inflammatory Bowel Disease, IBD). We conducted a LEfSe analyses [60] to unveil the key functional units/pathways that underlie the distinctions between the condition T2D vs. HHS and IBD vs. HHS.

Overall, our analysis identified a total of 17,225 KOs and 476 pathways across all samples. When applying the default parameters in LEfSe (log10 LDA > 2 and adjusted p-value < 0.05) to detect differentially abundant functional units, we identified 206 KOs and 71 pathways in the IBD group; 316 KOs and 83 pathways in the T2D group compared to the healthy control (HHS group). Figure 4 displays the top 10 results from the comparison between T2D and HSS. The most notable findings are functions related to glycan degradation (map00511) and sphingolipids (map00600), both of which have substantial supporting evidence for their involvement in disease status or development [11, 71, 79, 31, 76, 36, 1, 75, 26]. Besides, we further examined the results regarding [Carbohydrate](https://www.genome.jp/kegg/pathway.html#carbohydrate) [metabolism](https://www.genome.jp/kegg/pathway.html#carbohydrate), a well-established association in the context of type 2 diabetes [67]. Out of the 15 total pathways under this category, we found 9 significantly differential pathways between T2D and HSS samples (map00020, 00040, 00051, 00052, 00500, 00520, 00630, 00640, and 00660) though they are not the most significant hits, indicating a strong link to Carbohydrate metabolism.

**Figure 4:**
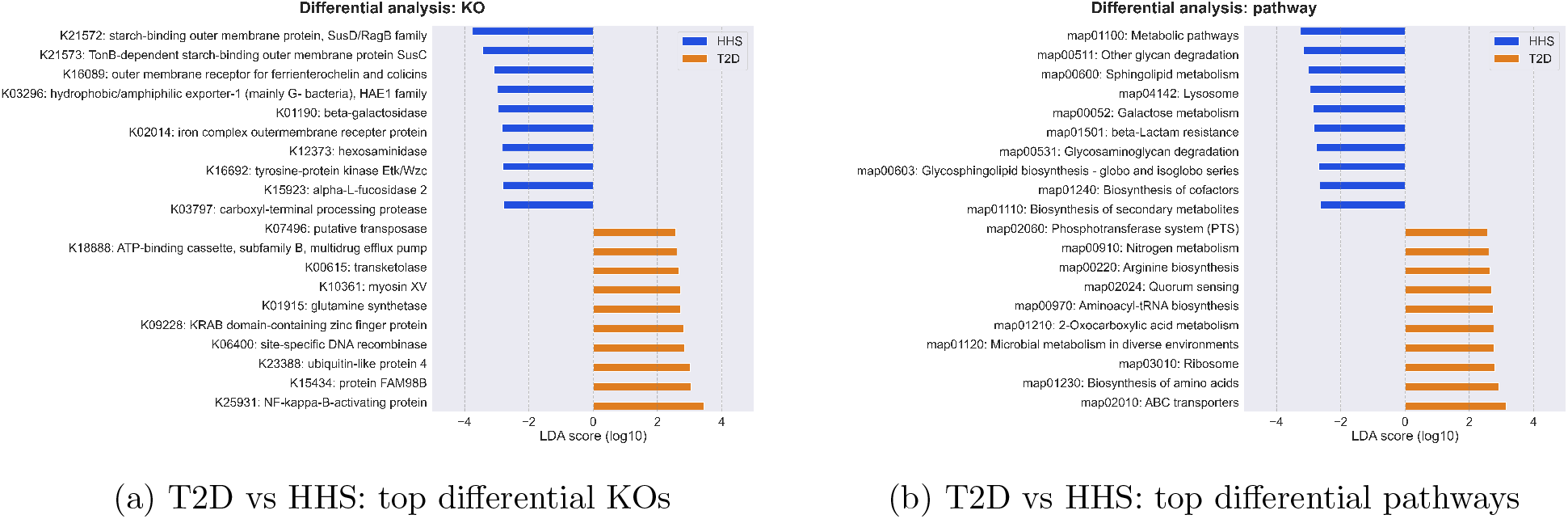
Differential analysis for T2D vs HSS. (a) Top 10 differential KOs in T2D samples compared to healthy samples. (b) Top 10 differential KEGG pathways in T2D samples compared to healthy samples.

**Figure 5:**
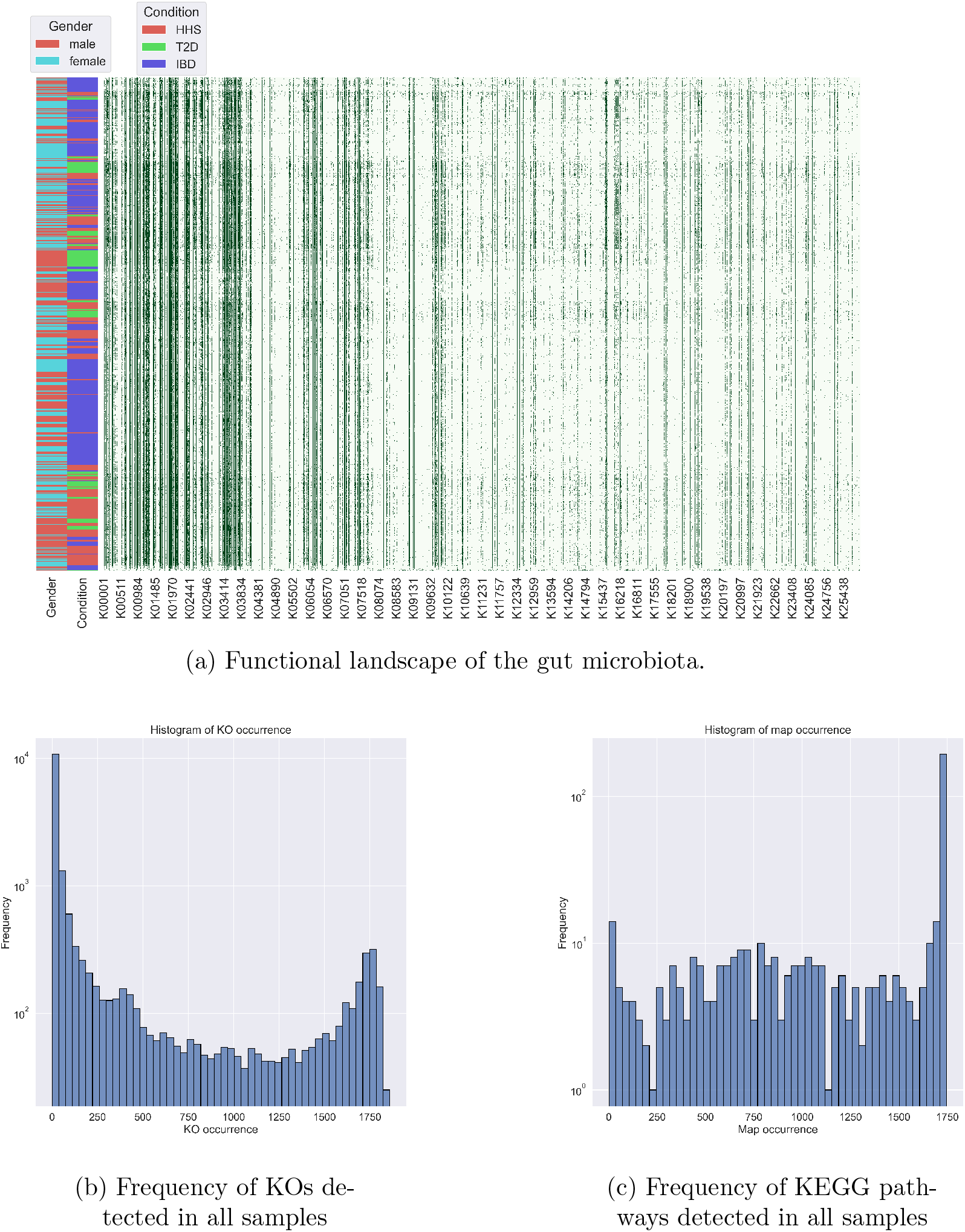
Functional landscape of the gut microbiota. (a) Heatmap of KO occurrences in all gut metagenomic samples from HMP database. Each row represents one metagenomic sample and each column shows the presence of a given KO across all samples. Samples are clustered by KO profiles. A vertical green “line” suggests that a KO shows up in most of the samples. (b) Frequency distribution of KO occurrences: the right side contains KOs identified in most of the samples, i.e. “housekeeping”. (c) Frequency distribution of KEGG pathway occurrence.

Digging into the KO-level analysis, the most significantly differentially abundant hit is NF-kappa-B-activating protein (K25931), which may play a key role in the pathogenesis of vascular complications of diabetes [65]. We also found starch-binding outer membrane protein (K21572), hydrophobic/amphiphilic exporter-1 (K03296), and carboxyl-terminal processing protease (K03797) etc. Though not illustrated here, we observe consistent functional signatures that corroborate existing knowledge from the comparisons between IBD and HHS groups as well. These observations substantiate the reliability of these functional profiles and may provide valuable insights for further investigating the functional roles of the gut microbiome. FracMinHash in conjunction with the KEGG database may serve as a fast and accurate hypothesis-generating methodology within the realm of functional analysis.

## 4 Methods

### 4.1 Overview of the pipeline

Traditional gene identification tools work on a given list of genes and assign sequencing reads in a metagenome to a list of potential genes using alignment. In contrast, as a *k*-mer-based approach, we found that we can decompose genes and sequencing reads (in the input metagenome) into sets of *k*-mers, and use `sourmash prefetch` to find genes present in the metagenome. `sourmash prefetch` takes the FracMinHash sketch of a query and finds overlaps with a list of reference sketches. The degree of overlap is determined by the Containment index, which is reported in the output.

Ideally, sketches of all genes can be used to obtain a functional profile of a given metagenome. The problem with using a list of genes in a sketching-themed setting is that gene lengths are too small for FracMinHash sketches to pick up a representative number of *k*-mers. Indeed, `sourmash` is intended to sketch many large sequence sets, not massive amounts of small sequences. On that account, although we took inspiration from an exhaustive list of genes, we abandoned this approach.

Therefore, instead of finding genes, we focus on finding the orthologous gene *groups* in a metagenome. We designed our pipeline `fmh-funprofiler` to detect gene groups as follows: `fmh-funprofiler` takes a list of orthologous gene groups as input, splits them into *k*-mers, and computes the FracMinHash sketches of these groups. These sketches are used as the reference sketches. The pipeline also takes a metagenome as input and obtains the FracMinHash sketch of the metagenome the same way. The pipeline then runs `sourmash prefetch` to find which gene groups have an overlap with the input metagenome. We designed `fmh-funprofiler` to invoke `prefetch` using the following parameters: `--protein scaled=1000 threshold bp=1000`. The scale factor 0.001 (equivalent to `scaled=1000`) is the default in `sourmash`. The parameter `threshold bp` controls when to stop searching for an overlap. We experimented with various values of `threshold bp` and found a good balance of speed and accuracy at 1000. The degree of overlap reported by `sourmash prefetch` is used to obtain the relative abundances of the orthologous gene groups in the metagenome.

As a proxy for the orthologous gene database, we picked the KEGG database in this work, which groups genes into orthologous groups called ‘KEGG Orthologs’ (referred to as KOs). The number of KOs (a total of only 25K) is much smaller than the number of genes, and the number of *k*-mers in a KO is much larger than that of a single gene – making the KOs an ideal choice for a sketching-based application. Additionally, the use of KEGG Orthology (KO) for functional comparison in metagenomics is a common practice for higher-level analysis[42, 35, 38], as evidenced by its application in tools like CAMISIM[15] and DIAMOND[8]. It is important to be aware that this method would miss genes that are not in the KO families.

### 4.2 Datasets

#### 4.2.1 Metagenome simulation

In this section, we describe how the simulated metagenomes were designed. There are many metagenome simulation tools available. We used CAMISIM [16] because of its extensive use in literature, and its ability to use a taxonomy – making it a “close to real-life” choice. We used CAMISIM [16] to simulate a metagenome from a random selection of genomes from all 4498 bacterial genomes present in the KEGG database, as of June 2023. Metagenomes can be complex and large [9], with the number of organisms varying from mere hundreds to many thousands. We ran CAMISIM to simulate metagenomes using a random selection of 64 genomes from all bacterial genomes. The actual selection is determined by the random seed fed to the CAMISIM program. This number 64 was chosen arbitrarily: we also checked for a range of other numbers of genomes and found similar results.

We can invoke CAMISIM using a number of read-simulators. For the analyses presented here, we used `wgsim`. with its default error profile. A mean fragment size of 270 was used with a standard deviation of 27. Other read-simulators or other choices of fragment mean and standard deviation do not affect the results significantly. CAMISIM selects the strains of the genomes using an input taxonomy. We used NCBI taxonomy for this step, setting one strain per OTU. And finally, CAMISIM chooses the relative abundances off of a Lognormal distribution. We used the default parameters (mean = 1 and standard deviation = 2) as parameters of this Lognormal distribution.

#### 4.2.2 Ground truth

To benchmark the performance of a tool on these simulated data, we needed to construct the following ground truth: which KOs are truly present in the metagenome, and what are their relative abundances? We construct this ground truth as follows: besides the sequencing reads, CAMISIM also outputs alignment mapping of all these reads. For every reference genome, we collected the start and end coordinates of all genes present in that reference genome from the KEGG database. We then parsed the alignment mapping files using the genes’ start and end coordinates to find the number of nucleotides of all *genes* present in the simulated metagenome. Next, we used the gene-to-KO mapping to calculate the total number of nucleotides of all *KOs* present in the metagenome. Finally, we used the number of covered nucleotides to calculate the relative abundances of the KOs.

#### 4.2.3 Reference data

In order to invoke our pipeline, we need FracMinHash sketches of all KOs. This section describes how these were obtained and constructed. The KEGG data, including KEGG Orthology (KO), gene sequences, and protein sequences), were downloaded from the KEGG FTP server, as of June 2023. We used the KO identifiers that belong to the KEGG brite hierarchy **Ko00001** which contains 25,413 KOs. After filtering out the KOs without protein sequences, we were left with a total of 25,347 KOs used for the downstream analysis. The hierarchical structure of **Ko00001** was used as “KEGG KO tree”.

We employed `sourmash` to compute FracMinHash sketches from amino acid sequences from those 25,347 KOs. These FracMinHash sketches were formulated using the command `sourmash sketch` with parameters `-p protein, k=7, k=11, k=15, abund, scaled=1000` (that is to say, we use *k*-mers for amino acid sequences and generate FracMinHash sketches using a scaling factor of 1000 for three different k values: 7, 11, and 15. Besides, *k*-mer abundances are tracked for quantification purposes). Subsequently, a sequence bloom tree (SBT) was constructed from the sketched KO database to accelerate the profiling process, though this is not required. These reference data are freely available on [Zenodo](https://zenodo.org/records/10045253).

#### 4.2.4 Real metagenomes

All human metagenomic samples were sourced from the HMP data portal [69], using specific search terms: “feces” for body sites, “fastq” for format, and “WGS raw seq set” for type. On January 2023, a total of 3550 files were retrieved, tagged under the manifest id “160bdc491e”. However, for quality assurance, additional filters were applied: (1) 185 files from the “MOMS-PI” project remained private; (2) 6 files were eliminated due to broken links; (3) 691 files exhibited unmatched MD5 values compared to metadata; (4) 4 files were corrupt; and (5) 148 files were identified as assembled scaffolds. Consequently, a final selection of 2516 files (a total of 4.6TB) was used. In subsequent analysis, we found numerous samples with very low sequencing depth. Therefore, we excluded files with sizes less than 200MB and files with less than 1000 KOs identified. After all these filtered steps, we have 1747 high-quality data remaining for the downstream analysis, including 547 healthy samples, 274 type 2 diabetes samples, and 926 samples related to inflammatory bowel disease.

## 5 Discussion

### 5.1 Conclusions

In this manuscript, we combined the FracMinHash sketching technique along with the KEGG orthology to construct a robust pipeline for generating functional profiles from metagenomic data. Our findings demonstrate that the functional profiles derived through this pipeline exhibit advantages in being a faster, more lightweight, and nearly as accurate alternative to `DIAMOND`, a widely used alignment-based tool for functional profiling. Naturally, if the resources allow for it, alignment-based results will be the most accurate, and alignment mapping information can reveal many biologically meaningful insights. However, the size of the data will not always remain manageable. When the data becomes too large for alignment-based tools, sketching-based tools can comfortably handle the data, probably without requiring a commercial workstation. While sketching will miss low abundance entities, the rate of false positives is low, and the results obtained on the detected entities appear trustworthy.

By using this pipeline, we conducted a comprehensive functional annotation analysis on all available human gut metagenomic data from the Human Microbiome Project[69], comprising a total of 2.5K samples (in which 1747 samples were used for downstream analysis). We successfully generated sample-specific functional signatures and performed differential analyses across different conditions. The top signatures we identified align with existing literature. Moreover, the KO-based functional profiles expanded our insights by introducing additional disease-related functional units. We believe this pipeline holds significant value in the realm of metagenomic functional profiles, especially when scalability and hypothesis-generating analyses are paramount.

### 5.2 Going beyond functional profiles

Gene Ontology (GO) enrichment analysis is widely used for understanding biological processes in experimental datasets. KEGG is such a comprehensive database that offers structured modules and pathways that delineate functional elements, associations with diseases, and interactions with drugs. It serves as an invaluable resource for exploration and hypothesis generation in diverse research endeavors. The potential of this direction can be fully exploited through the integration of functional profiles with meticulously curated biomedical knowledge graphs (e.g. Hetionet [24], RTX-KG2 [74]). This integration represents an exciting avenue for advancing our comprehension of intricate biological systems: metagenomic functional profiles provide insights into the functional potential of microbial communities; KEGG’s rich resource of hierarchical modules and pathways offers a structured framework to interpret and contextualize these functions; and knowledge graphs, which link genes, proteins, diseases, drugs, and other biological entities, can facilitate the discovery of underlying relationships between different entities. By connecting functional profiles with biomedical knowledge graphs, researchers can explore the functional intersections of diseases [57, 51], identify potential therapeutic targets [78, 81, 48], and predict novel associations between microbial functions and human health [17, 46]. Pursuing this integrated approach can not only aid in hypothesis generation but also pave the way for innovative research.

## A Definitions of metrics used

In this section, we describe the metrics we used to benchmark `fmh-funprofiler` and `DIAMOND`. We first give a high-level textual definition, and then give the rigorous mathematical formula. To write the mathematical formulas, we need to introduce a few notations first.

Let the set of KOs truly present in the ground truth be 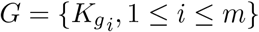, and the set of KOs predicted by a tool be 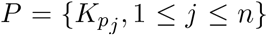. Also, let the abundance of a KO 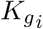 in the ground truth be 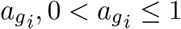, and the abundance of a KO 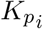 in the predicted output be 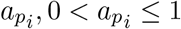. We also assume that the KOs in the ground truth are ordered based on the non-increasing order of abundances, i.e. for any 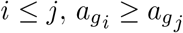.

The metrics we used in our analyses are:

- Precision: the fraction of KOs that are truly present amongst the discovered KOs.

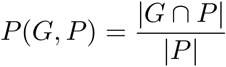
- Completeness: the fraction of the gold standard KOs identified by the tools.

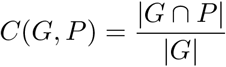
- Completeness in top 95% KOs: the fraction of the top 95% abundant gold standard KOs identified by the tools. To mathematically define this, let 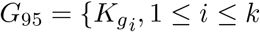, where *k* is the smallest integer that 0.95}. Then, this metric is as follows:

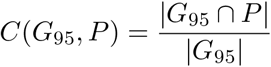
- Weighted Jaccard similarity: weighted Jaccard index of the abundances of the KOs in the ground truth, and those identified by the tools. To define this, we define the weight functions *w*_*G*_ and *w*_*P*_ as follows: if *K* ∈ *G*, then *w*_*G*_(*K*) gives the abundance of *K* in ground truth, and 0 otherwise. Also, if *K* ∈ *P*, then *w*_*P*_ (*K*) gives the abundance of *K* in the prediction, and 0 otherwise. Using these functions, weighted Jaccard similarity is defined as follows:

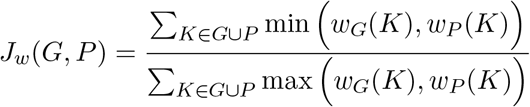
- Correlation: the Pearson correlation coefficient of the abundances of the ground truth KOs, and those identified by the tools (only limited to the correctly identified KOs). Standard Pearson correlation was used, and therefore, we are not including the mathematical definition for this metric here.
- Bray-Curtis distance: the Bray-Curtis distance of the abundances of the ground truth KOs, and those identified by the tools (only limited to the correctly identified KOs). Bra-Curtis distance is often used to measure the dissimilarity in species composition between two samples. We used Bray-Curtis to quantify the dissimilarity in KO composition.

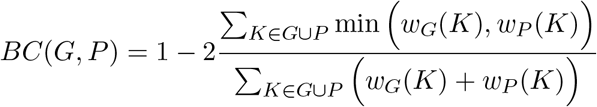

## B Functional landscape of the gut microbiota

## C Details of functional profiles of HMP data

We used command `sourmash sketch` with parameters `-p protein, k=11, abund, scaled=1000` for FracMinHash sketches of the HMP data (though it’s suggested for the reference data to have multiple *k* values to fit various purposes, one proper *k* size is sufficient for functional profiles). Next, functional profiles of them were generated using `sourmash prefetch` with parameters `-k` `11 --protein --threshold-bp 500` (*k* = 11, which corresponds to 33 in DNA sequences, is a reasonable value for protein sequence comparison).

## D Details of the differential analysis presented in Section 4

LEfSe (Linear discriminant analysis Effect Size) was performed to determine enrichment in functional profiles of pairs of conditions. We adhered to the default settings for this analysis. Features (KO or pathway) with LDA score (log10) higher than 2 and adjusted p-value < 0.05 were recognized as significant. We used the Seaborn package in Python for plotting. For simplicity, we only showed partial results in the manuscript.

